# Parasitoid wasp response to infection with a reproductive manipulator symbiont reveals subtle changes in host proteome in life-long symbiosis

**DOI:** 10.64898/2026.11.24.768972

**Authors:** Olivia L. Mathieson, Suzanne E. Kelly, Matthew R. Doremus, Dylan L. Schultz, Stephan Schmitz-Esser, Martha S. Hunter, Manuel Kleiner

## Abstract

*Encarsia* is a genus of minute parasitoid wasp of the family Aphelinidae. Interestingly, many species of this genus carry bacterial symbionts, such as the reproductive manipulator *Cardinium*. Most of what is known about *Encarsia* physiology comes from studies on their ecology, morphology, and reproductive phenotypes associated with symbiont interactions, however, there is limited research exploring the molecular physiology of these species and the broader Aphelinidae family. Here, we present proteomes of male and female adult *Encarsia suzannae* with and without the cytoplasmic incompatibility inducing symbiont *Cardinium hertigii c*Eper1. We identified proteins that respond to symbiont infection and propose potential targets for symbiont-host cell manipulation and reproductive phenotypes. We also characterized the expressed innate immune capacity of the wasp and found that expression patterns were suggestive of a limited immune response toward *Cardinium*. Lastly, we identified expression differences in proteins putatively involved in sensory biology which may provide insights into cryptic *Cardinium*-induced phenotypes. Overall, our data suggest that subtle differences in *E. suzannae* physiology may support life-long *Cardinium* association. These data provide a molecular overview of *E. suzannae* and can be used as the basis for further characterization of *Encarsia* and, more broadly, a reference for the molecular physiology of Aphelinidae.

## Introduction

Inherited (or vertically transmitted) bacterial symbionts are widespread across insects (Moran et al., 2008). Many of these symbionts, which are intracellular and obligately associated with their hosts, are passed from mother to offspring directly packaged inside the egg (transovarial transmission), and thus fully reliant on their hosts to complete their lifecycle (Perlmutter and Bordenstein, 2020). Symbiont induced host reproductive manipulations, phenotypes in which symbiont produced effectors directly modulate host reproduction to favor symbiont transmission from one generation to the next, may have evolved out of this pressure (Doremus and Hunter, 2020; Duron et al., 2008; Werren et al., 2008). These symbiont-induced phenotypes are impactful from an evolutionary perspective: “sabotaging” uninfected offspring and occasionally skewing sex ratios in the host population to promote symbiont spread (Doremus and Hunter, 2020) and possibly contributing to host speciation (Bruzzese et al., 2022; Gebiola et al., 2016). Likely due to the impact of reproductive phenotypes, most of the research on reproductive manipulator symbionts focuses on the mechanisms underlying these phenotypes. Some of these symbionts, however, are found in somatic cells beyond the reproductive organs, as is the case for the Bacteroidota symbiont *Cardinium* in its host *Encarsia suzannae* (Mathieson et al., 2026), yet the systemic effects of these symbionts are commonly underexplored.

*Encarsia* is a genus of minute parasitoid wasp from the family Aphelinidae. Parasitoid wasps use different insect species to complete their lifecycle, often laying their eggs inside host insect species (Peters et al., 2017). As a result of this life history characteristic, parasitoids in the family Aphelinidae play an ecological role in controlling populations of plant-feeding insects such as whitefly, scale insects, and aphids (Hunter and Woolley, 2001; Polaszek et al., 2026).

With the natural host range of these wasps overlapping with common agricultural pest species, it is not surprising that aphelinid wasps are employed in pest management efforts. In fact, import and targeted release of *Encarsia* and other aphelinids have been used historically to control invasive whitefly populations in the United States (Ciomperlik and Goolsby, 2008; Gould et al., 2008; Nguyen and Bennett, 1995). Much of the work on *Encarsia* centers on just a few species. One of these is *Encarsia suzannae*, an aphelinid parasitoid of *Bemisia tabaci* (whitefly) that was originally collected in Weslaco, Texas, USA (Gebiola et al., 2017b). While much research has been conducted on the biology of this wasp (Donnell and Hunter, 2002), there is limited molecular data available for this species (Schultz et al., 2022) and thus characterization of the molecular underpinnings of *E. suzannae* immunology, sensory biology, reproductive biology, and metabolism remain vastly underexplored.

Most of what is known about *E. suzannae* focuses on its association with the transovarially transmitted intracellular reproductive manipulator symbiont *Cardinium* (Hunter et al., 2003; Zchori-Fein et al., 2004). *Cardinium* induces a cytoplasmic incompatibility (CI) phenotype in *E. suzannae* that prevents proper development and survival of embryos produced from symbiont-carrying males mated with symbiont-free females (Gebiola et al., 2017a; Hunter et al., 2003). Research on *E. suzannae-Cardinium* interactions has focused on symbiont localization in reproductive organs, influence on host behavior, the molecular biology of the symbiont, and the CI phenotype (Doremus et al., 2020, 2019; Gebiola et al., 2017a; Harris et al., 2010; Mann et al., 2017; Mathieson et al., 2026; Penz et al., 2012; Perlman et al., 2014; Schultz et al., 2025), however, many questions remain about this symbiosis regarding the mechanism of the CI phenotype and, generally, interactions between the symbiont and somatic host cells.

Several studies have sought to explore more general *Cardinium-*host interactions in *B. tabaci* MED (whitefly) and *Dermatophagoides farinae* (mites) through bulk -omics methods (Hubert et al., 2021a, 2021b; Li et al., 2018a, 2018b; Liu et al., 2023; Yang et al., 2021, 2023) or non-native host cell culture (Nakamura et al., 2011), though studies exploring hosts with CI-inducing *Cardinium* strains or considering fundamental aspects of host biology such as sex are needed (Mathieson et al., 2025). With *Cardinium* being a widespread symbiont across multiple arthropod and non-arthropod invertebrate hosts (Mathieson et al., 2025), understanding *Cardinium* host-interactions in diverse host groups is critical. Specifically, understanding how symbionts, like *Cardinium*, interact with biocontrol species such as *E. suzannae* may provide insights into not only population dynamics and drivers of wasp evolution, but also molecular physiology of the wasps and how that ultimately influences their ecological role and effectiveness in pest management.

Several aphelinid species commonly carry inherited bacterial symbionts that, in the case of *E. suzannae*, directly influence host reproduction. Yet very little is known about interactions between the wasps and their symbionts beyond the important reproductive phenotypes. To better understand the molecular physiology of *E. suzannae* and its interactions with its symbiont *Cardinium*, we analyzed the proteome of adult male and female *E. suzannae* with and without (generationally cured) *Cardinium*. We found limited proteomic response to *Cardinium* and thus propose that *E. suzannae* may subtly shift metabolism to compensate for lifelong *Cardinium* infection by changing expression of specific metabolic proteins, rather than changing entire pathways. Additionally, we identified several putative cell cycle or mitotic proteins that respond to symbiont infection suggesting possible systemic interactions between *Cardinium* and the *E. suzannae* cell cycle, offering possible targets for symbiont-induced CI. We also identified proteins putatively involved in the *E. suzannae* immune system, including several proteins across multiple innate immune categories that are sex-specific. Lastly, we identified expressed odorant binding proteins (OBPs) and found that several exhibit sex- and symbiosis-specific expression patterns. Combined, our results provide critical insights into the molecular physiology of *E. suzannae* and its interactions with its symbiont, bringing to light many unknowns regarding how *Cardinium* may influence its hosts.

## Results and Discussion

### Proteome Overview

We measured the proteome of adult male and female *E. suzannae* with and without the reproductive manipulator symbiont *Cardinium hertigii c*Eper1. To increase the depth of our proteomic measurement, we subjected pooled 100-wasp homogenates to a differential centrifugation fractionation approach which produced three distinct proteomic fractions (**Fig. 1a**) (Hinzke et al., 2018; Mathieson et al., 2026). Through this, we detected 10,423 *E. suzannae* proteins and 8,864 were quantified via area under the curve (AUC) quantification, ranging from approximately 2,500 to 3,500 proteins per sample per fraction on average (**Supplementary Fig. S1**, **Supplementary Data 1**). Of these, 81.4% were assigned to orthogroups inferred by OrthoFinder (Emms and Kelly, 2019) with default parameters using the following proteomes as reference: *Eretmocerus hayati*, *Nasonia vitripennis*, *Trichogramma brassicae*, *Apis mellifera*, *Bemisia tabaci*, *Galleria mellonella*, *Bombyx mori*, and *Drosophila melanogaster*. There were 483 proteins belonging to *E. suzannae*-specific orthogroups and 1,943 unassigned proteins also unique to *E. suzannae*. We also detected 235 proteins of the *C. hertigii c*Eper1 symbiont, which were removed from downstream analyses.

**Figure 1.**
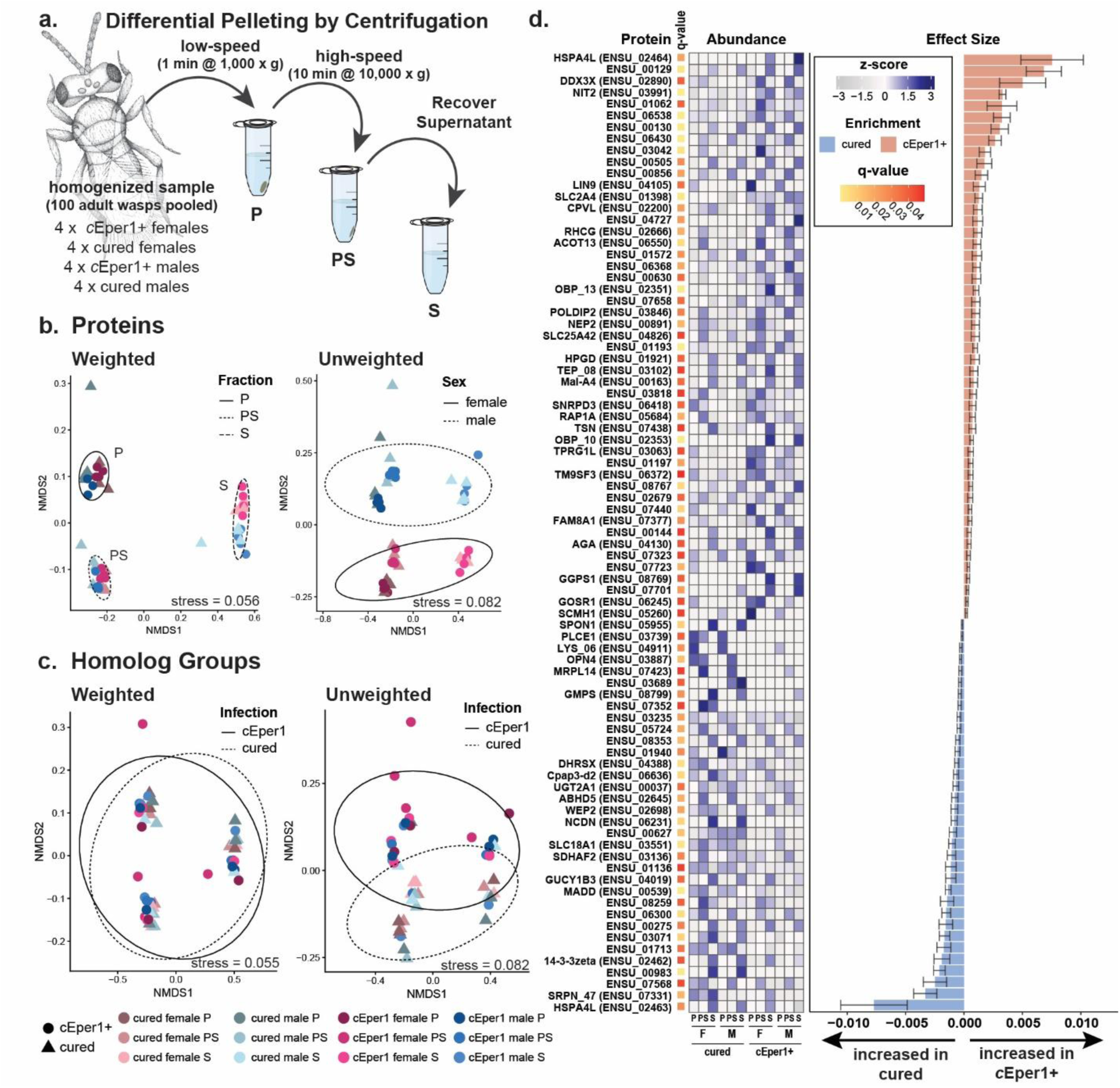
Overview of the *Encarsia suzannae* proteome. (a) Experimental overview depicting sample types used in this study and the fractionation method used to increase proteome depth. (b) Non-metric multidimensional scaling plots showing proteomic profile at the protein level. Each point represents a single sample proteome, color of the points indicates sample type (fraction, sex, infection status), and shape indicates infection status. The left plot shows weighted distances (Bray-Curtis) and ellipses show fraction clustering at 90% confidence. The right plot shows unweighted distances (Jaccard) and ellipses show clustering by sex at 90% confidence. (c) Non-metric multidimensional scaling plots showing proteome composition summed by orthogroup (see methods). Each point represents a single sample proteome based on abundance profiles of orthogroups, and color and shape of point indicate sample type and infection status. The left plot shows weighted distances (Bray-Curtis) and the right plot shows unweighted distances (Jaccard). In both plots, ellipses show infection status clustering at 90% confidence. (d) Differentially expressed proteins between cured and symbiont-carrying wasps (MaAsLin2, q < 0.05). Heatmap depicts averaged relative abundance (TSS-AUC) square root transformed and z-scored by row. The bar plot shows effect size of infection with error bars indicating standard error associated with the model.

Proteome composition was predominantly affected by fraction (P, PS, & S) at the protein level, with each fraction having a distinct compositional profile (**Fig. 1b**, **Supplementary Data 2**; p-value_fraction_= 0.001, adonis 2, Bray-Curtis). Differences in these fractions were also observed by fluorescence imaging (**Supplementary Fig. S2**). The P fraction, or the low-speed pellet, was most heterogeneous containing large intact pieces of cuticle, antennae, and wings and other autofluorescent components of the host. Large DAPI foci in this fraction are likely cell nuclei.

Supporting this, the P fraction is enriched in GO terms associated with larger structures, such as chromatin and nuclear-localized biological processes (e.g., nucleus organization, regulation of transcription), muscle fibers, and cellular junctions (**Supplementary Fig. S3**, **Supplementary Data 3,4**). The PS fraction, or high-speed pellet, contained an enrichment of *c*Eper1 cells and DAPI foci of varying morphologies. For example, smaller DAPI puncta may be mitochondria while elongated DAPI signals are likely sperm cells, in addition to some nucleus-like puncta (**Supplementary Fig. S2**). Accordingly, the PS fraction proteome was enriched in mitochondrial and other organelle proteins (e.g., organellar ribosomes) (**Supplementary Fig. S3**, **Supplementary Data 3,4**). The S fraction, or final supernatant, contained limited fluorescence signal, consistent with it containing extracellular or non-DNA carrying components of the cytoplasm and extracellular fluids (e.g., hemolymph) (**Supplementary Fig. S2**). The S fraction proteome was enriched in the extracellular and cytosolic metabolic components of the proteome as expected (**Supplementary Fig. S3**, **Supplementary Data 3,4**). Microscopy, GO enrichment analysis, and similar compositional dispersion of the proteomes across fractions suggest that these fractions produce consistent proteomic profiles that each provide distinct and relevant biological information (**Supplementary Table S1**).

At the protein level, separation between the sexes was more distinct when we used an unweighted distance metric (**Fig. 1b**), suggesting low abundance proteins, likely sex-specific isoforms, may be driving differences in proteomes between sexes. The influence of infection status on proteomes was masked by the larger differences of fraction and sex. When we summed protein abundances by orthogroup, the influence of fraction and sex on proteome composition was reduced, while the influence of infection became clearer (**Fig. 1c**). The loss of separation between fractions and sexes supports our idea that summing by orthogroup likely merged sex-specific, tissue-specific, and cellular compartment-specific isoforms of proteins into single units averaging out specific signals associated with these factors. Interestingly, the infection-based separation appears more distinct when using an unweighted distance metric, though not statistically significant. This suggests that the compositional differences at the orthogroup-level may also be driven by lower abundance groups.

To determine the effects of *Cardinium* infection on expression of *E. suzannae* proteins, we compared the proteomes of symbiont-carrying and symbiont-free adult wasps at the protein and orthogroup level. We found a moderate response to *Cardinium* in the *E. suzannae* proteome, with 83 proteins and 39 orthogroups significantly responding to symbiont infection (**Fig. 1d, 2a**, **Supplementary Data 5,6**). The majority of the responding proteins can be grouped into three main functional categories: metabolism, cell growth and proliferation, and signaling. In addition there are several proteins involved in intracellular transport and phagocytosis, as well as putative venom proteases and many proteins of unknown function with significant responses to *c*Eper1 infection (**Fig. 1d**).

### Expression of immune-related proteins suggests a sexually dimorphic immune system and limited immune response to the symbiont

To understand the immune capacity of *E. suzannae*, we mined protein annotations to identify immune proteins. In total, we identified 245 proteins likely involved in *E*. *suzannae* immune function from 23 previously described groups of innate immune proteins (**Fig. 3**, **Supplementary Fig. S4**, **Supplementary Data 7,8**; Waterhouse et al., 2007). Of these proteins, 215 have at least one predicted domain supporting homology-based immune protein group assignment. Further, 22 immune proteins were unique to *E. suzannae* based on orthogroup assignment. Serpins (SRPN) were the largest immune category with 65 proteins identified (**Fig. 3a**), followed by C-type lectins (CTL) with 35 proteins. Other detected categories included peroxidases, with thioredoxins (TPX) being the largest group, scavenger receptors (SCR), inhibitors of apoptosis (IAP), CLIP-domain serine proteases (CLIP), and (pro)phenoloxidases (PPO) **(Fig. 3a**). We identified proteins from several immune groups with sex-specific detection patterns. In most of these cases, expression was female-specific. This is strongly demonstrated in the 16 female-specific SRPNs (**Fig. 3b**). Other examples of sex-specific immune expression include female-specific toll-like receptors (TOLLs) and PPOs as well as female- and male-specific CTLs and MD2-like proteins (ML) (**Fig. 3b**). Caspases (CASPs), like CASP_02, were also more consistently detected in females compared to males.

We also relied on detection patterns of immune proteins across sample fractions to provide insights into potential functional roles of the predicted immune proteins. Many of the immune proteins were detected consistently (i.e., in nearly all replicates) across all fractions (**Fig. 3b**, **Supplementary Fig. S4**). However, when fraction-specific detection of immune proteins was observed, it was predominantly in the S fraction where extracellular and cytosolic components were enriched (**Supplementary Fig. S3**). For example, MLs, CTLs, TOLLs, and peptidoglycan recognition proteins (PGRP) proteins exhibited an S fraction-specific detection pattern (**Fig. 3b**, **Supplementary Fig. S4**). A similar pattern was also observed in some CLIPs, lysozymes (LYS), and SRPNs (**Fig. 3b**). Overall, immune system proteins did not appear to respond to *Cardinium* infection, although in some cases they were reduced in abundance.

We observed sexual dimorphism across several innate immune protein groups in *E. suzannae* with females having more sex-specific immune proteins. This is consistent with observations of sexual dimorphism of the immune system across the animal kingdom (Belmonte et al., 2020; Kelly et al., 2018). Additionally, many innate immune proteins share homology with proteins involved in development. It has been proposed that innate immunity arose out of the need to distinguish between self and malignant self at the basis of multicellular life (Mushegian and Medzhitov, 2001). For this reason, many innate immune proteins likely have context-dependent functions and are still involved in central developmental pathways, such as metamorphosis and embryogenesis (Darin et al., 2025; Sun et al., 2004), limiting the extent to which we can characterize such proteins based on sequence similarity alone.

Evolution and functional co-opting of immune-related proteins is not only a remnant of the large landmark evolutionary transitions, but it is an ongoing evolutionary process. For example, annotated immune proteins with a primary function of controlling the immune response have undergone expansion events that have potentially allowed for emergence of novel functions at the organismal level. These proteins can be co-opted for interactions with other species, such as anticoagulants in the saliva of blood feeding insects and protease cocktails in venoms (Meekins et al., 2017). Parasitoid wasps, like *E. suzannae*, are reliant on interactions with host insects to complete their reproductive cycles and must combat host defense mechanism (e.g., melanization) against the parasitoid egg (Reed et al., 2007; Sullivan and Neuenschwander, 1988). For example, central inhibitors of immune cascades, like SRPNs, have been shown to be included in parasitoid venoms to directly inhibit activation of these cascades in the host insect in response to damage caused by the oviposition of the parasitoid egg itself (Colinet et al., 2013, 2009; Meekins et al., 2017). It is possible that some of the predicted female-specific SRPNs observed in this study function in a similar manner and should be investigated further as potential components of the wasp venom.

Direction of expression response of certain immune proteins to symbiont infection may be indicative of tolerance between *E. suzannae* and *c*Eper1. This is surprising, given that *Cardinium* is generally proposed to be more immunogenic than other symbionts, especially when compared to the distantly related reproductive manipulator *Wolbachia*. For example, *Cardinium* infection was shown to upregulate several immune genes in silkworm cell culture, such as a peptidoglycan recognition protein gene and antibacterial protein genes (e.g., *lysozyme*, *lebocin 4*, *cecropin B*) (Nakamura et al., 2011). *Cardinium* has even been hypothesized to contribute to the allergenicity associated with dust mites (Yang et al., 2023). We, however, did not observe similar immunogenic responses at the proteome level, suggesting a degree of host- or symbiont-mediated immunological tolerance toward the symbiont.

In support of symbiont tolerance, we observed decreased expression of lysozyme-like LYS_06 and some serpins (SRPN_47 and SRPN_14) in symbiont carrying wasps (**Fig. 3d-e**). Lysozymes are antibacterial enzymes that directly inhibit bacterial growth through degrading the peptidoglycan in bacterial cell walls. Unlike some other bacterial symbionts which have lost their capacity to synthesize peptidoglycan (Otten et al., 2018), *Cardinium* retains most of its peptidoglycan biosynthesis pathways and likely its peptidoglycan layer (Kurtti et al., 1996; Penz et al., 2012). LYS_06 shares homology with other lysozyme-like proteins, however, it does not contain lysozyme-specific functional domains. Instead, LYS_06 has a “prokaryotic membrane lipoprotein lipid attachment site profile”, a protein domain previously associated with eukaryotic antibacterial defensins (Islam et al., 2023). This suggests a role for this protein in microbial interaction, and absence of it in symbiont-carrying wasps may support the idea of immune tolerance towards *Cardinium* in this host. Similarly, some SRPNs are decreased in response to symbiont infection. Serpins, which are regulators of protease signaling cascades, have been previously shown to be upregulated in response to bacteria (Eappen et al., 2013; Woon Shin et al., 1998). Therefore, reduction could indicate reduced immune capacity during symbiont infection, perhaps to tolerate the symbiont.

Another interesting infection-based difference is the increased expression of CASP_02 in infected females (**Fig. 3c**). Increased CASP expression has been previously associated with *Wolbachia*, a distantly related reproductive manipulator endosymbiont, where it is proposed to increase rates of oogenesis through promoting apoptotic turnover of follicle cells in ovaries of infected hosts (Guo et al., 2018). It is possible that *Cardinium* relies on a similar mechanism to promote host oogenesis and, consequently, its own transmission. While these ideas are intriguing, it is important to consider that a large portion of the innate immune system is post-translationally regulated (e.g., phosphorylation and protease cascades), so studies that go beyond proteomics, along with deeper annotation efforts and functional characterization of proteins, will be necessary to understand the full impact of *Cardinium* on the *E. suzannae* immune system.

### *Encarsia* metabolic response to *Cardinium* infection suggests possible mechanisms for metabolic compensation and symbiont control

Our results suggest that *E. suzannae* proteomic response to *Cardinium* seems to be largely driven by changes in a small subset of proteins of diverse functions, rather than changes in full pathways. For example, overall cell metabolism appears consistent in the two groups (**Supplementary Fig. S5**), yet specific proteins involved in cell metabolism significantly respond to *c*Eper1 infection (**Fig. 1d, Supplementary Fig. S6**). These include proteins involved in energy production, such as the increased expression of mitochondrial complex I assembly factor (ENSU_06368), a NADH:ubiquinone oxidoreductase subunit (ENSU_06430), and phosphoglycerate mutase (ENSU_02679) (**Supplementary Fig. S6**). Complementing this, we observed increased expression in response to symbiont infection of some cellular transporters (e.g., ABC transporters, sugar transporters, ammonium transporters; **Supplementary Fig. S6**) and several proteases and peptidases (ENSU_07701, ENSU_00891 (Nep2), ENSU_02200 (CPVL); **Fig. 1d**). This response also included changes in homologs of proteins involved in production of oxidative compounds and/or balancing host cell metabolism under oxidative stress, such as a decrease in an oxidoreductase (ENSU_00983) as well as increased expression of a carboxylesterase (ENSU_00129) and an AMP-binding luciferase-like enzyme (ENSU_03042) (**Fig. 1d**). Other proteins that decreased in response to symbiont infection were guanine monophosphate synthetase (GMPS, ENSU_08799) and succinate dehydrogenase (SDH) assembly factor 2 (ENSU_03136). In addition, these proteins and other differentially expressed proteins are potentially involved in anabolic pathways, such as fatty acid biosynthesis, amino acid biosynthesis, and nucleic acid biosynthesis. Notably, we did not observe large abundance changes of proteins associated with these pathways responding in unison to symbiont infection, rather changes were more variable and were not consistently in a single direction (i.e., up or down in response to infection; **Supplementary Fig. S5**).

The diverse host responses to *Cardinium* infection being driven by specific proteins rather than entire pathway responses is similar to previous work demonstrating mixed correlations of *Cardinium* and mite host transcripts (Hubert et al., 2021a) and functionally diverse proteomic responses to *Cardinium* in whitefly (Li et al., 2018a). However, both of these studies suggested a stronger differential expression response to *Cardinium* than we observed. Further, work identifying miRNA with target functions of the miRNAs ranging from molecular and cellular functions to broader systemic organismal functions responding to *Cardinium* infection in whitefly (Li et al., 2018b) also supports the functionally diverse response we observed. We propose that these diverse, and in our case limited, responses are likely reflective of the life-long, symbiotic relationship of *Cardinium* and its host in contrast to what we might expect from acute pathogenic infection. Alternatively, because this study is comparing infected to cured wasp line, not a never-infected wasp line, this limited response could reflect that many of the major changes that we might expect in response to infection are an evolved phenotype that is retained even in the absence of the symbiont.

Unlike pathogens, it is thought that symbionts generally do not influence host translation and instead may employ effectors that post-translationally modify host proteins to “curate” their cellular environment (Ribet and Cossart, 2010; Russell and Castillo, 2020). In support of this, *Cardinium* has been previously associated with changes in acetylation of host histones, proteins involved in metabolic processes, and transporters in whiteflies (Li et al., 2018a). While symbiont effectors could be mediating these effects, it is possible that specific changes in expression of host-derived modification enzymes may drive host protein modifications. For example, we observed changes in expression of specific host metabolic enzymes that may be involved in post-translational modifications and secondary signaling metabolite production, like omega-amidase, glycosyltransferase, and glycosyl hydrolase (**Fig. 1**), which might suggest altered patterns of post translational modifications of *E. suzannae* proteins associated with *Cardinium* infection.

Though the overall response is limited, specific proteins responding to symbiont infection may suggest subtle rewiring of the host cell metabolism to compensate for increased metabolic burden associated with life-long symbiont infection, as seen in other long-term symbiont infections (Jacobs et al., 2026). Like many other obligate intracellular symbionts, *Cardinium* has limited metabolic capacity and thus relies on its host to meet most metabolic requirements, like amino acids, nucleic acids, and other essential molecules (Penz et al., 2012; Schultz et al., 2025). Therefore, increases in specific metabolic enzymes and some transporters in the *E. suzannae* proteome may suggest increased metabolic substrate availability to compensate for the nutritional requirements of *Cardinium*. We observed increased expression of an omega-amidase, which may suggest elevated production of certain amino acids, such as glutamine and asparagine (Cooper et al., 2016). In insects, this protein is broadly uncharacterized, though proposed to be involved in functions from development and metabolism (Pletcher et al., 2019; Rotelli et al., 2019) to insecticide resistance (Liu et al., 2020), suggesting the metabolic role of omega-amidase homologs in insects may be co-opted for a larger diversity of functions in a species-specific context. We also observed an increase in several putative proteases (e.g., ENSU_07701, ENSU_00891 (Nep2), ENSU_02200 (CPVL)). Increased expression of some proteases could provide an example of how *Cardinium* may restructure host processes to meet nutritional requirements. For example, intracellular pathogens are known to upregulate misfolded protein stress response in host cells to take advantage of nutritional byproducts of host catabolic processes (Cornejo et al., 2017; Russell and Castillo, 2020). Perhaps the observed increased expression of some proteases, and possible subsequent cleaved amino acids, serves as an amino acid source for the symbiont.

We also observed an increase in some proteins involved in fatty acid and lipid metabolism (e.g., enoyl-(acyl carrier protein) reductases, polyprenyl synthetase, fatty acyl-CoA reductase), and a reduction of proteins for using possible products of this metabolism for energy production (e.g., Cytochrome P450) (**Supplementary Fig. S7**). Alteration of lipid metabolism has broadly been shown in response to infection with bacterial pathogens and symbionts in insects (Visser and Scheifler, 2026). Altered lipid metabolism has been attributed to a range of symbiont functions from dependence on host lipids for symbiont proliferation to maintaining and promoting host lipid metabolism to induce protective effects through symbiont competition for lipids with host antagonists and vectored diseases (Attardo et al., 2023; Herren et al., 2014; Lindsey et al., 2018; Paredes et al., 2016; Shigenobu et al., 2026). In *Encarsia*, variable differences observed in lipid metabolism-related proteins may suggest that lipid production and/or production of fatty acid-based secondary metabolites (e.g., steroids, hormones) may be affected by *Cardinium* infection. As with amino acids, this could suggest that *Cardinium* infection modifies host fatty acid metabolism to acquire necessary fatty acids and lipid precursors for its own growth.

Alternatively, it is possible that host lipid production could increase to accommodate altered endoplasmic reticulum (ER) and Golgi body restructuring. In support of this idea, we observed increased expression of tSNARE (ENSU_06245), a central component of membrane organization in and trafficking between the ER and Golgi body, during *Cardinium* infection (**Fig. 1d, 2c**). Intracellular bacterial pathogens and symbionts, like *Wolbachia*, have been shown to restructure and use host cell membranous organelles (e.g., ER and Golgi body) to escape host immune recognition and concentrate nutritional requirements to promote proliferation (Fattouh et al., 2019; Gurumurthy et al., 2014; Heuer et al., 2009; Russell and Castillo, 2020). While observations of *Cardinium* within host cells are limited, *Cardinium* from *Ixodes scapularis* (tick) have been reported to influence the structure intracellular membranes (e.g., vacuolation, formation of lysosomes and other membrane vesicles) of infected cells in culture (Kurtti et al., 1996). This previous report combined with our proteomic observation suggests that *Cardinium* may restructure membranous intracellular host cell structures like other symbionts and intracellular pathogens, though follow-up studies are needed. Ultimately, the combination of increased and decreased expression of metabolic proteins potentially reflects balancing of symbiont and host nutrient demands, specific alignment of host and symbiont physiology, and, possibly, symbiont-mediated remodeling of the host cell.

**Figure 2.**
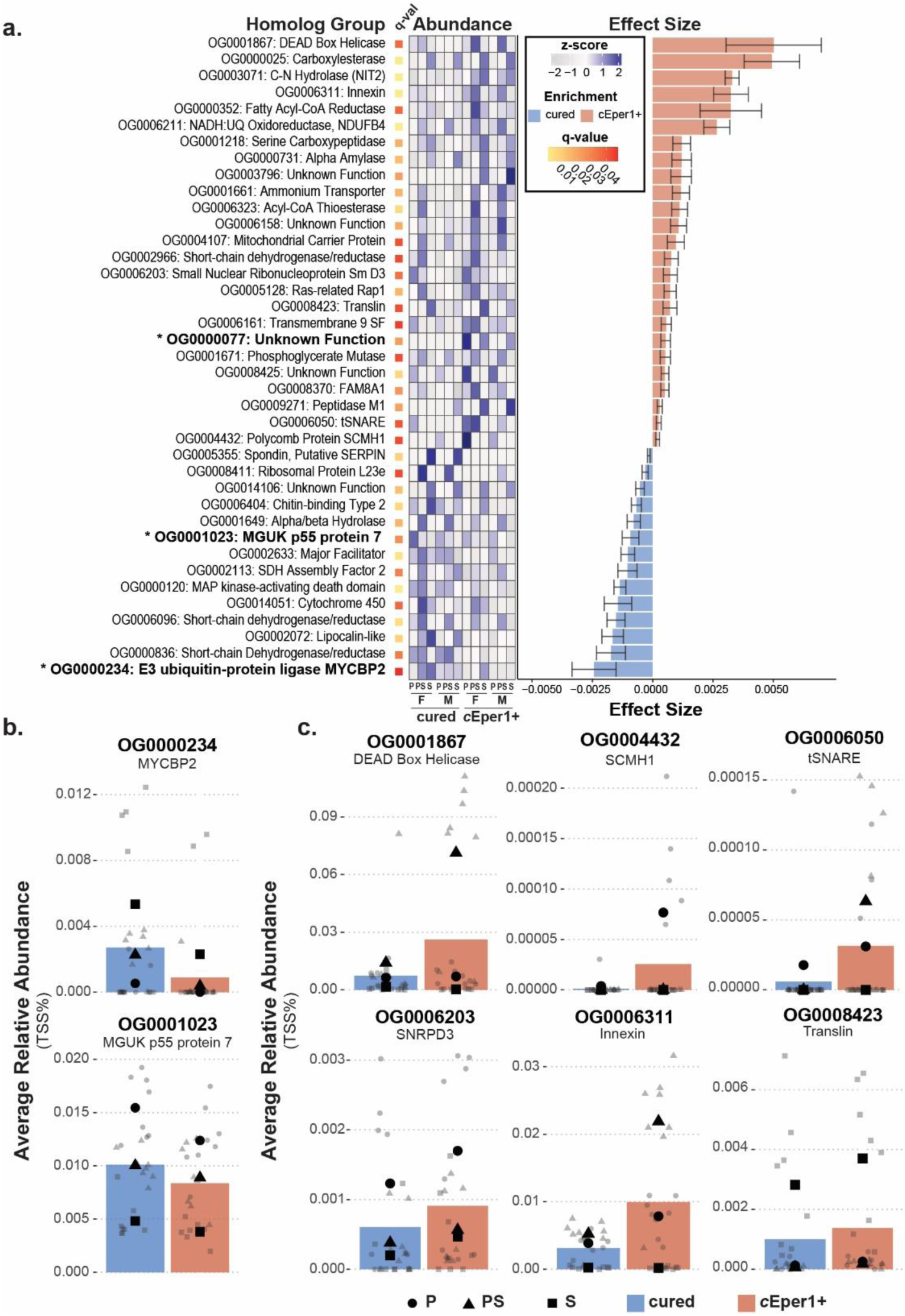
Differential expression of *E. suzannae* orthogroups, highlighting proteins putatively involved in signaling and regulation. (a) Differentially expressed homolog groups between cured and symbiont-carrying wasps (MaAsLin2, q < 0.05). Heatmap depicts averaged summed relative abundance (TSS-AUC) of homolog groups square root transformed and z-scored. The bar plot shows effect size of infection response with error bars indicating standard error associated with the model. Bolded homolog groups highlight groups that were not deemed significant in the protein-level differential expression analysis (see Figure 1). (b) Regulatory and signaling orthogroups significantly decreased in response to symbiont infection. (c) Regulatory and signaling orthogroups significantly increased in response to symbiont infection. For both (b) and (c), bars represent average summed relative abundance (TSS-AUC) on a percent scale for a single orthogroup by infection status, each point represents summed relative abundance of proteins in the orthogroup per replicate, and shape indicate proteome fraction. Larger black shapes indicate the average of the fraction (circle = P fraction, triangle = PS fraction, square = S fraction). Colors: blue = cured, orange = *c*Eper1+.

**Figure 3.**
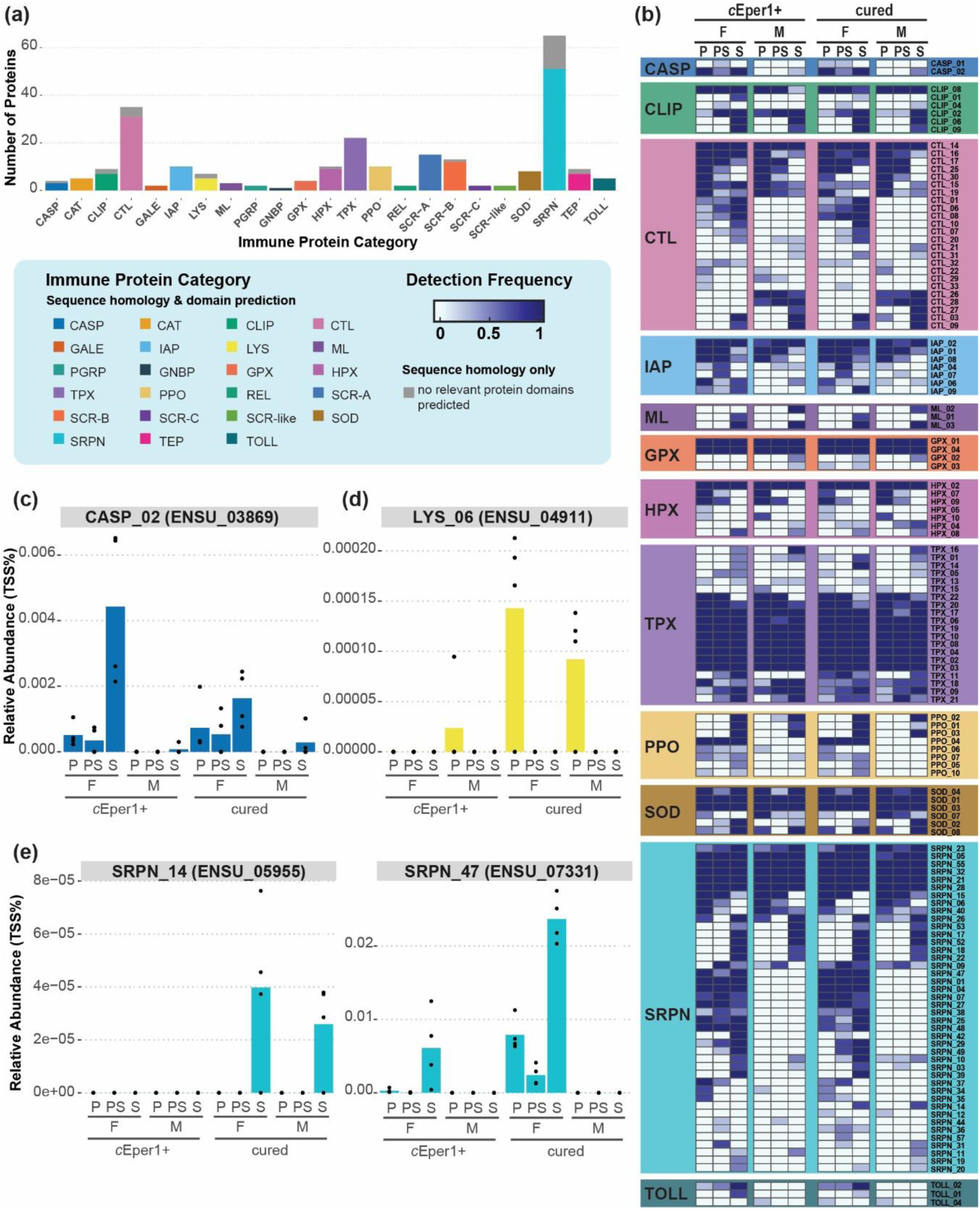
Proteomic detection of the *Encarsia suzannae* immune system. (a) AUC-quantifiable proteins putatively involved in the *E. suzannae* immune system identified through sequence homology and inclusive domain prediction (i.e., must have at least one relevant domain). (b) Frequency of detection of immune proteins by category across fraction, sex, and infection status. (c-e) Relative abundance of CASP_02, LYS_06, SRPN_14, and SRPN_47 across fraction, sex, and infection status. Bars indicate mean abundance and each point indicates a single sample. Abbreviations: CASP = caspases, CAT = catalases, CLIP = CLIP-domain serine proteases, CTL = c-type lectins, GALE = galectins, IAP = inhibitors of apoptosis, LYS = lysozymes, ML = MD2-like proteins, PGRP = peptidoglycan recognition proteins, GNBP = Gram-negative binding proteins, GPX = glutathione peroxidases, HPX = heme peroxidases, TPX = thioredoxin peroxidases, PPO = (pro)phenoloxidase, REL = Rel-like NFkappaB proteins, SCR-A (B, C, like) = scavenger receptors class A (B,C, like), SOD = superoxide dismutases, SRPN = serine protease inhibitors, TEP = thio-ester containing proteins, TOLL = toll-like receptors.

### *Cardinium* infection is associated with changes in proteins involved in cell proliferation

Several proteins potentially involved in cell growth and proliferation respond to symbiont infection. Proteins that were increased in symbiont-carrying wasps include nuclear protein MDM1 (ENSU_01136), polycomb protein SCMH1 (ENSU_05260), LIN9 protein (ENSU_04105), innexin (ENSU_6538), small nuclear ribonucleoprotein Sm D3 (SNRPD3; ENSU_06418), translin (ENSU_07438), and DEAD box helicase (ENSU_02890) homologs (**Fig. 1d & 2b,c**, **Supplementary Data 9**). In contrast, we observed reduction of protein orthogroups potentially involved in signaling, such as MAP kinase-activating death domain (OG0000120), MYCBP2 (OG0000234), and MAGUK p55 protein 7 (OG0001023), in *c*Eper1-carrying wasps (**Fig. 2c**). Combined, these results suggest that *Cardinium* may influence cell division and stress signaling pathways.

The *c*Eper1 *Cardinium* strain induces cytoplasmic incompatibility (CI) in *E. suzannae* (Hunter et al., 2003). CI is a phenotype that results in failed embryogenesis of offspring produced when a symbiont-carrying male and a symbiont-free female mate. In *E. suzannae*, CI ultimately causes embryonic death following multiple repeated attempts at cell division during embryonic development in which chromosomes do not fully separate during mitosis resulting in an amorphous mass of cell-like structures and clouds of chromatin (Gebiola et al., 2017a). This is similar to the cytological defects induced by the more widely studied CI-inducing *Wolbachia* (Callaini et al., 1997, 1996; Tram et al., 2006). The proposed mechanism and impacts of the CI phenotype strongly suggest that *Cardinium* produces effectors that interact with the cell cycle, perhaps through structurally interfering with mechanisms of mitosis and/or regulation of cell cycle checkpoints. Even though the CI phenotype occurs during early embryonic development, it is possible that these effectors act beyond this life stage.

One of the two *Cardinium* effectors proposed to induce CI in *E. suzannae*, CAHE_p0043, leads to diffuse nuclear morphology, DNA damage response, and failed cell division when the effector is expressed in yeast (Mathieson et al., 2026). We observed decreased expression of a nuclear protein MDM1 homolog, a protein that binds microtubules and is a negative regulator of centriole duplication (Van de Mark et al., 2015), along with increased expression of several proteins that share homology with proteins involved in chromatin structure and cell proliferation in the context of eukaryotic models (**Supplementary Data 9**; Aoki et al., 1995; Benton and Johnston, 2003; Esterlechner et al., 2013; Heard, 2005; Hong et al., 2022; Koedoot et al., 2021; Martinez and Cavalli, 2006; Neff et al., 2023; Sahni et al., 2017; Schievella et al., 1997; Stucke et al., 2007; Tsuda et al., 2019; Wang et al., 2004; Xu et al., 2018). These proteins could be potential targets of *Cardinium*-induced CI. Beyond CI, altered expression of proteins involved in chromatin structure, cell growth and proliferation in the bulk proteome of *c*Eper1-carrying wasps could suggest integration of timing between *Cardinium* proliferation and host cell growth and regulation. These mechanisms would most likely be host- and strain-specific. Combined these results suggest that *Cardinium* may interact with host cell division processes systemically, in addition to its CI-related cell division impacts.

### Changes in odorant binding protein expression in response to *Cardinium* may provide insights into cryptic symbiont-induced host phenotypes

Several odorant binding proteins (OBPs) and other putative sensory proteins were among the proteins responding to symbiont infection. These proteins mostly decreased in abundance, except for two OBPs, OBP_10 and OBP_13, which were consistently detected in only *c*Eper1-carrying wasps (qvalue_OBP_10_ = 0.0002, MaAsLin2; qvalue_OBP_13_ = 0.0005, MaAsLin2; **Fig. 1d, 4b**). We explored this further through annotating additional OBPs, identifying 50 OBPs in the *E. suzannae* proteome (**Fig. 4**, **Supplementary Data 10**). Most of these were orthologs of known OBPs from other insect groups, however, we identified 4 OBPs that appear to be unique to *E. suzannae* (OBP_04, OBP_38, OBP_47, and OBP_49). Most OBPs were detected exclusively in the S fraction, suggesting possible circulation in the hemolymph; however, a small portion of OBPs were detected across fractions (**Fig. 4a**). Of these, 10 OBPs exhibited sex-specific expression, with 9 out of the 10 being specific to females.

**Figure 4.**
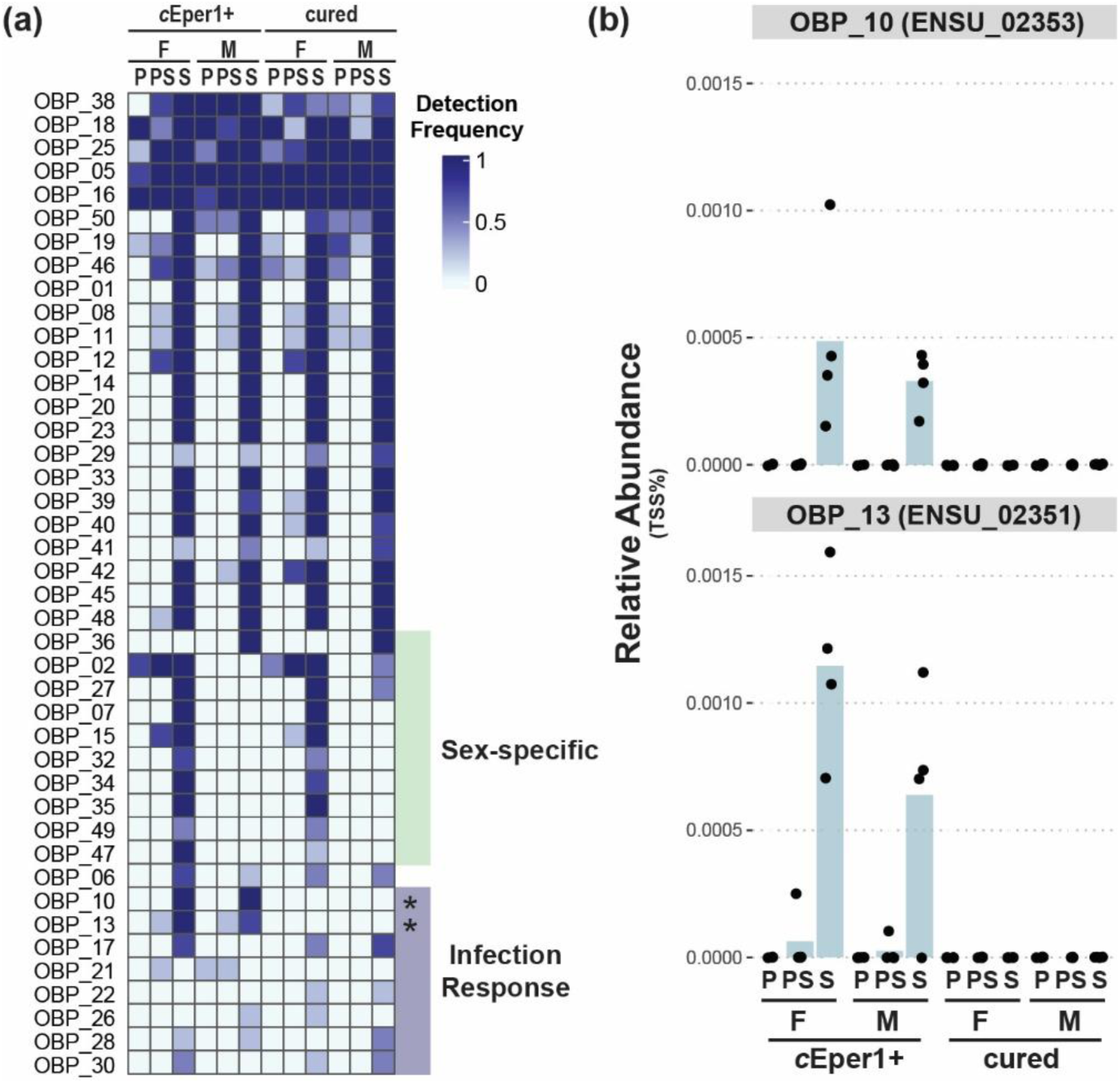
Proteomic detection of *Encarsia suzannae* odorant binding proteins. (a) Frequency of detection of *E. suzannae* odorant binding proteins (OBPs) across fraction, sex, and symbiont status. Included are OBPs detected at least twice in the full dataset. Green shading indicates sex-specific detection of OBPs and purple shading indicates infection responding OBPs. Cured versus *c*Eper1+ differential expression significance indicated by ‘*’ (MaAsLin2; q < 0.2). (b) Relative abundance of OBP_10 and OBP_13 showing symbiosis-specific expression in both male and female *E. suzannae*.

Specific expression of OBPs in symbiont-carrying wasps suggests a role for these OBPs in symbiont recognition, interaction, or symbiosis-associated sensory phenotypes. OBPs are central to insect physiology through sensory, development, and immune signaling (Abendroth et al., 2023). In parasitoid wasps, OBPs have been suggested to be involved in probing behavior, or host selection (Zhu et al., 2024). While there is little to no information available on *Encarsia* OBPs, studies conducted in other *Encarsia*-*Cardinium* symbioses demonstrate that *Cardinium* may influence host oviposition behavior (Kenyon and Hunter, 2007; Zchori-Fein et al., 2001). While behavior modifications were not observed in *c*Eper1-carrying *E. suzannae* in this study (Kenyon and Hunter, 2007), it is possible that *Cardinium* may subtly alter wasp host choice or other behaviors not measurable in the lab.

Additionally, observation of symbiosis-specific expression of OBPs is interesting within the context of *Cardinium* transmission dynamics at the population level. In a population cage study, CI *Cardinium* spread more rapidly than predicted from CI strength and fecundity costs alone, suggesting that *Cardinium* may induce cryptic beneficial effects in the host promoting its spread (Harris et al., 2010). It is possible the observed changes in OBP profiles may provide the molecular underpinnings of cryptic fitness effects. Future work focused on understanding how these putative differences in sensory biology may influence reproductive physiology and symbiont spread dynamics in more complex populations, where both symbiont-carrying and symbiont-free wasps are interacting, will provide insights into the behavior phenotypes associated with these molecular findings.

### Limitations and Future Priorities

Our results provide critical insights into the molecular physiology of *E. suzannae* and its potential interactions with its reproductive manipulator *Cardinium*. While depth of measurement was comprehensive through subsampling by fractionation, this was a bulk proteomic measurement, and most cell type- and tissue-specific physiological signals were likely lost.

Ultimately, this is likely why we did not observe clear full pathway responses to the symbiont in this host. Tissue-resolved or single-cell proteomics and/or transcriptomics provide a promising avenue to continue exploring the impacts of *Cardinium* on host molecular physiology at a higher resolution. Because of the complexity of protein function within a cell and, more broadly, a multicellular organism (Sigaeva et al., 2026), it is difficult to assign function to proteins in a non-model system where molecular measurements and the capacity for sufficient follow-up for these measurements is limited. Advances that have allowed for work on obligate intracellular bacteria like *Wolbachia* and *Rickettsia* in cell culture have provided critical insights into bacteria-host cell interactions and broader cellular physiology (reviewed: Porter and Sullivan, 2023; Sit and Lamason, 2024) and recent molecular modification advances have opened up interrogation capacity of these traditionally challenging systems (Lamason et al., 2018; McGinn et al., 2024; Nadal-Jimenez et al., 2019; Penunuri et al., 2026). Adopting similar approaches in the *Encarsia-Cardinium* system would be necessary to interrogate many of the emerging functional hypotheses put forth in this study.

Aside from host-symbiont interactions, our orthogroup analysis suggests many homologs are present within the *E. suzannae* proteome. It is important to consider that many of these are likely isoforms, however, these observations could also suggest gene duplication events. Genomic sequencing of *E. suzannae* and other *Encarsia* species will be necessary for proper identification of paralogs and, subsequently, clearer distinction between isoforms and gene duplication events. Identifying gene duplication events and overall patterns of evolution of genes involved in sensory biology, immunology, and secondary metabolism are an important part of *Encarsia* biology to explore and may be critical for informing biocontrol efforts involving *Encarsia* and the broader Aphilinidae family (Polaszek et al., 2026).

## Conclusion

We measured the proteome of adult male and female *E. suzannae*, a parasitoid wasp of the family Aphelinidae, with and without its reproductive manipulator symbiont *Cardinium* to explore the interactions between host and symbiont. We found that *E. suzannae* has a diverse functional response to *Cardinium* infection at the protein-level, with variable metabolic, cell signaling, and proliferation responses. We also identified *E. suzannae* immune proteins, several of which exhibited sex-specific expression patterns, but showed a limited overall immune response to the symbiont. Lastly, we identified two putative OBPs that were specific to symbiont carrying wasps suggesting phenotypic impacts of the symbiosis may go beyond reproductive manipulation. Combined, these results provide a first look into the molecular physiology of *Encarsia* and the integration of the physiologies of *E. suzannae* and *Cardinium*, expanding our understanding of this ecologically and economically relevant insect and its relationship with its reproductive manipulator symbiont.

## Materials and Methods

### Insect rearing and experimental design

We used cultures of *E. suzannae*, host of *Cardinium c*Eper1, maintained at the University of Arizona in Tucson, Arizona, USA for this study. These cultures were originally collected from South Texas, USA and have since been maintained at 27°C with a 16h:8h day:night light cycle. Female *E. suzannae* are raised on *Bemisia tabaci* (whiteflies) infested cowpea (*Vigna unguiculata*) and males, which are hyperparasitoids, are raised on whiteflies parasitized by *Eretmocerus emiratus* (Hunter et al., 2003). Cured lines of *E. suzannae* were established by treating symbiont carrying wasps with rifampicin for 3 generations and were maintained for greater than 10 generations prior to experimental use (Hunter et al., 2003). For this study, we produced four replicates each of symbiont carrying adult males, symbiont carrying adult females, cured adult males, and cured adult females with each replicate consisting of 100 wasps pooled to obtain enough biomass for metaproteomic measurement. Due to the high number of wasps, we reared wasps in batches over the course of a month and blocked the study by replicate (i.e., we processed one replicate of each experimental group per batch) to prevent the experimental variables of sex and infection status from being confounded by batch effect (Wagner and Kleiner, 2025).

### Proteomic sample preparation

For each replicate, we thoroughly homogenized 100 wasps in 1 ml of 1X phosphate buffered saline (PBS) with protease inhibitor cocktail (1 tablet per 10 ml of PBS; cOmplete, Mini, EDTA-free Protease Inhibitor Cocktail, Roche: 04693159001) using a glass homogenizer (KIMBLE DUALL Tissue Grinder, DWK Life Sciences). We used a differential sedimentation rate centrifugation protocol to obtain three fractions from homogenates (Hinzke et al., 2018; Mathieson et al., 2026). Homogenates were centrifuged at 1,000 x g for 1 minute. The resulting loose pellet was retained as the first fraction (P fraction). The supernatant was carefully removed and subjected to further centrifugation at 10,000 x g for 10 minutes, with the resulting tight pellet retained as the second fraction (PS fraction) and the supernatant retained as the third fraction (S fraction). The S fraction was lyophilized for 15 hours to remove excess liquid and concentrate the samples. We suspended P and PS fractions in 60 μl of SDT lysis buffer (4% (w/v) SDS, 100 mM Tris-HCl pH 7.6, 0.1 M DTT) and the S fraction in 120 ul of SDT lysis buffer. Samples were heated at 95°C for 10 minutes and then cooled to room temperature. We then used a modified filter-aided sample preparation method to prepare samples for LC-MS/MS analysis (Hamann et al., 2016; Kriete et al., 2026; Wiśniewski, 2019). For this, cooled lysates were loaded onto centrifugal filters (Millipore, MPE010025) in 1 (P and PS fractions) or 2-4 (S fraction) loading steps. Each loading step loads 60 μl of lysate with 400 ul of urea buffer (8 M urea in 0.1 M Tris/HCl pH 8.5) and centrifuging at 14,000 x g for 40 minutes. After loading, filter units were washed with 200 μl of urea buffer and centrifuged at 14,000 x g for 20 minutes. Samples were then treated with 100 μl of IAA solution (0.05 M iodoacetamide, 8 M urea in 0.1 M Tris/HCl pH 8.5), mixed at 600 rpm for 1 minute in thermo mixer, allowed to incubate without mixing for 20 minutes, and then centrifuged at 14,000 x g for 20 minutes. We then washed filters with 3 urea buffer washes, each by adding 100 μl of urea buffer to the filter unit and centrifuging for 20 minutes at 14,000 x g. A buffer exchange to ABC (50 mM Ammonium Bicarbonate) was performed by adding 200 μl of ABC to filter units and centrifuging at 14,000 x g for 20 minutes. To remove excess hydrophobic residues from the filter unit, we washed filter units twice by adding 200 μl of ACN wash solution (60% ACN, 0.2% formic acid), mixing at 600 rpm for 1 minute in a thermos mixer, and then centrifuging at 14,000 x g for 20 minutes. We then further washed the filters with 3 ABC wash steps, each by adding 100 μl of ABC to the filter unit and centrifuging for 20 minutes at 14,000 x g. We transferred the filter units to new collection tubes. We added 0.8 μg of trypsin (Pierce Trypsin protease, MS grade) in 40 μl of ABC, mixed at 600 rpm for 1 minute, and then incubated in a wet chamber at 37°C for 12 hrs to allow for digestion. After incubation, filter units were centrifuged at 14,000 x g for 20 minutes to collect peptides. Lastly, 50 μl of 0.5 M NaCl with 0.4% formic acid was added to the filter units, mixed at 600 rpm for 1 minute in a thermo mixer, and centrifuged for 20 minutes are 14,000 x g to elute remaining peptides. We retained the eluate for LC-MS/MS analysis. We used the Pierce mBCA assay (Thermo Scientific) to determine peptide concentrations according to manufacturer instructions.

To visually confirm the content of fractions from the fractionation protocol, we fractionated as before and fixed samples of fractions to glass coverslips for fluorescent *in situ* hybridization (FISH) microscopy with a previously described method (Daims et al., 2004; Doremus et al., 2020). We fixed the homogenates by adding 4% paraformaldehyde in 1X PBS (PFA) to the sample on the slide and incubating for 20 minutes at room temperature. We then removed the PFA and rinsed with DI H_2_O for minute twice. The samples were dehydrated with subsequent 3-minute washes with 50%, 80%, and 96% EtOH and then allowed to dry for approximately 5 minutes. Once dry, we added 20 ul of hybridization buffer (10% formamide, 0.9 M NaCl, 2 mM Tris HCl pH8, 0.01% SDS) with 2 μl of cy3-labelled probe (30 ng/ul; previously published in Doremus et al., 2020), placed the coverslip in a hybridization chamber, and incubated in the dark for 2 hrs at 46°C. After incubation, the coverslip was removed from the hybridization chamber and placed in wash buffer (20 mM Tris HCl pH 8, 0.45 M NaCl) which was preheated to 48°C. The coverslip in wash buffer was placed into a water bath to maintain 48°C for 12 minutes. After, we removed the cover slip from the wash buffer and dipped (2 seconds) the cover slip in ice cold DI H_2_O. Coverslips were air dried before adding 20 μl of DAPI (10 μg/mL in 1X Tris-buffered saline with Tween-20 (TBST)) to each sample and incubating in the dark for 5 minutes. We rinsed the coverslips 3 times with 1X PBS for 1 minute each. The cover slip was mounted on the slide using mounting media (80% glycerol, 20% TBST with 2% *n*-propyl-gallate). We collected images using an Olympus BX53 compound microscope with TRITC (Brightline TRITC-A-Basic-ZERO, Semrock) and DAPI (Brightline DAPI-11608-OFF, Semrock) filters for visualizing the cy3 probe and DAPI stain, respectively. Images were taken at 1000x magnification under immersion oil. Due to variability in the FISH technique, we used images for qualitative comparison only and no quantitative analyses were performed on images. Raw images were edited in ImageJ. Raw images are presented if informative, and ImageJ was only used to add scale bars for these images. Contrast and brightness were adjusted for the following images to remove background: male S fraction (DAPI, TRITC), female S fraction (DAPI), male PS fraction (TRITC), and female PS fraction (TRITC). All adjustments were made evenly across the full image, when performed.

### LC-MS/MS, database construction, and protein quantification

We measure peptide using 1D-LC-MS/MS methods previously described (Kriete et al., 2026). Briefly, we loaded 500 ng of peptide onto a trap column (PepMap Neo Trap Cartridge, PN 174500; Thermo Scientific) using a VanquishNeo liquid chromatograph (Thermo Scientific). Peptides were then eluted onto a 75 cm Easy-Spray HPLC column (PN ES905, Thermo Scientific) for separation using a 70 min reverse phase gradient at 60°C. Eluting peptides were ionized via electrospray ionization with an EASY-Spray source prior to measurement by data dependent acquisition (DDA) with an Orbitrap ASTRAL mass spectrometer (Thermo Scientific). Our DDA method acquired precursor scans in the Orbitrap mass analyzer using a scan range of 380-1,400 m/z at 240,000 resolution with an AGC target of 10 million ions and a 200 ms injection time. Precursors were acquired using a 1 s cycle time with the most abundant ions being selected for fragmentation with a normalized collision energy of 32% by HCD. After selection, peptides were excluded from fragmentation for a dynamic exclusion duration of 10 seconds. We acquired MS2 scans for fragmented ions in the ASTRAL mass analyzer with a scan range of 150-2,000 m/z, AGC target of 5,000, injection time of 3 ms, and an isolation window of 1 m/z.

We built a custom database consisting of protein sequences from *c*Eper1 proteome (UP000003996), *E. suzannae* transcriptome (Schultz et al., 2022), *Bemisia tabaci* (host of *E. suzannae*; GCF_001854935.1) and its bacterial symbionts (*Hamiltonella*, UP000216438; *Portiera*, UP000215611; *Rickettsia*, UP000215668), and potential laboratory contaminants (Mellacheruvu et al., 2013) following current recommendations for database construction (Blakeley-Ruiz and Kleiner, 2022). We used our custom database and Proteome Discoverer Software version 2.3 (Thermo Scientific) to analyze our raw spectral data using a previously described method (Kriete et al., 2026). Briefly: We denoised the ASTRAL MS2 data using the Top N Peaks Filter node, retaining only the top 10 peaks in each 100 Da window. The Sequest HT node was used to search spectra using the following parameters: Trypsin (Full) with a maximum of 2 missed cleavages, 6 to 144 amino acid peptide length, 10 ppm precursor mass tolerance, 0.05 Da fragment mass tolerance, and a maximum of 3 equal dynamic modification (oxidation +15.995 Da [M], deamidation +0.984 Da [N, Q, R], N-terminus acetyl group +42.011 Da) allowed per peptide. We included a static carbamidomethylation (+57.021 Da [C]) modification in the method, as well. The Percolator node calculated peptide FDR confidence and we only retained peptide spectral matches (PSMs) with FDR < 0.05. We inferred proteins using the Protein FDR Validator node and classified proteins as high (protein FDR < 0.01) and medium (0.01 < FDR < 0.05) confidence. We used an area under the curve (AUC) method to quantify the proteins with the Minora Feature Detector node using a minimum chromatographic trace length of 5 non-zero points and a PSM high confidence. We used the Feature Mapper node with ‘perform RT alignment’ set to false. We determined protein abundance using the Precursor Ions Quantifier node using ‘unique + razor’ peptides, considering protein groups for uniqueness, and calculating protein abundance based on summed peptide abundance. We exported individual results files from ProteomeDiscoverer and compiled them using a custom python script which retained only proteins that had at least medium-level protein FDR confidence, were classified as ‘master proteins’, and could be quantified using the AUC method.

### Statistical and functional data analysis

We shortened protein accessions (derived from transcript identifiers previously published by Schultz et al., 2022) by assigning protein identifiers with the following naming scheme: “ENSU_” for *<u>En</u>carsia <u>su</u>zannae* followed by a sequentially assigned number (00001 to 10423).

For proteins with manually curated protein names and EggNOG mapper-assigned protein names, we preferentially used these descriptions in figures and describing results. We used EggNOG Mapper (version emapper-2.1.13; Buchfink et al., 2021; Cantalapiedra et al., 2021; Huerta-Cepas et al., 2019) and InterProScan (version 5.59-91.0; Blum et al., 2021; Jones et al., 2014) via the Galaxy server (The Galaxy Community, 2026) to predict COG, GO, KEGG, PFAM, and other annotation information for detected *E. suzannae* proteins. We also identified putative homologs of *E. suzannae* proteins in available UniProt reference proteomes for well annotated insect models (*Drosophila melanogaster*, UP000000803; *Bombyx mori*, UP000005204; *Galleria mellonella*, UP001652740; *Apis mellifera*, UP000005203) and other parasitoid wasps (*Eretmocerus hayati*, UP001239111; *Nasonia vitripennis*, UP000002358; *Trichogramma brassicae*, UP000479190) using OrthoFinder (version 2.5.5; Emms and Kelly, 2019) through the Galaxy server (The Galaxy Community, 2026). OrthoFinder assigns groupings of protein homologs (inclusive of paralogs and orthologs) referred to as “orthogroups”. For *E. suzannae* proteins that were placed in orthogroups with proteins from the insect reference proteomes, we used the reference proteome protein descriptions as an extension of the annotations curated for the *E. suzannae* proteins. All annotation data is available in **Supplementary Data 11.**

We used functional annotation data of detected proteins to characterize innate immune proteins and odorant binding proteins. We developed an iterative key term search and refinement workflow to mine proteins putatively belonging to various immune protein categories associated with the insect innate immune system (Waterhouse et al., 2007). Immune proteins that passed the initial search and refinement steps were scored as such: 0 = immune protein category assignment is based on homology alone, 1 = immune protein category assignment is supported by prediction of at least one relevant functional domain. Immune and odorant binding proteins were assigned “curated protein names” using the naming scheme: immune protein abbreviation (Waterhouse et al., 2007) or “OBP” for “odorant binding protein” followed by a sequential number. These curated identifiers were preferentially used over the “ENSU_” protein identifiers and the EggNOG mapper-assigned protein names when describing these proteins in the study. Specific key terms for search and refinement of each protein group are available **Supplementary Data 7.**

For all statistical analysis, we analyzed two levels of data: the protein-level and orthogroup-level (i.e., summed protein abundances with proteins not assigned to orthogroups removed from the analysis). AUC quantification data was filtered to remove symbiont and contaminant proteins prior to normalizing the data via total sum scaling (TSS). For compositional analyses, we performed a square root transformation on TSS normalized data. We then performed nonmetric multidimensional scaling on Bray-Curtis dissimilarity and Jaccard index distances using metaMDS from the R package vegan (version 2.6-4) to visualize similarity of proteome composition across sample types (Oksanen et al., 2026). We assessed statistical significance of clustering patterns using adonis2 and betadisper from the R package vegan (version 2.6-4) (Oksanen et al., 2026).

To assess functional enrichment in the proteomic fractions, we performed a pairwise Welch’s T-test on filtered, normalized, transformed, and imputed data to compare each fraction to the others (e.g., P fraction vs. not P fraction). Briefly, TSS normalized data was filtered to retain only proteins present in at least one third of the dataset (i.e., 16 out 48 samples) and remaining missing values were imputed with a constant that was half the lowest value in the data frame before log2 transformation. We connected the Gene Ontology (GO) terms assigned via EggNOGMapper and InterProScan to the coreGO ontology (go.obo; Ashburner et al., 2000; The Gene Ontology Consortium, 2026) using the ontologyIndex (version 2.12) R package. We used the R package clusterProfiler (version 4.6.2) to assess GO term enrichment in each fraction against the total proteome with an over representation analysis using the enricher function (Xu et al., 2024; Yu et al., 2012).

For differential expression analyses, we followed recommendations laid out by Hinzke et al. (2026). Missing values were imputed with zero before the data were square root transformed. We used MaAsLin2 (version 1.12.0) to test for differential expression via linear mixed effects model (analysis_method = “LM”) with a Benjamini-Hochberg FDR p-value adjustment (Mallick et al., 2021). We allowed for random effects associated with homogenate replicate and used the following fixed effects: sex, infection status, and fraction.

All analyses were performed using R (version 4.2.2) and full statistical results are available in **Supplementary Data 2-6**. We used the following R packages for data cleaning prior to all analyses: ontologyIndex, tidyverse, dplyr, broom, tidyr, and stringr. We used the following R packages for data visualization: ggplot2 (version 4.0.2), ggforce (version 0.4.2), ggpubr (version 0.6.3), ggplotify (version 0.1.3), ggfortify (version 0.4.19), gridExtra (version 2.3), cowplot (version 1.2.0), ComplexHeatmap (version 2.14.0), and RColorBrewer (version 1.1-3). We adapted code, in part, from code generated using ChatGPT, Google Gemini, and various collaborative programming blogs for the purpose of data wrangling, high throughput code adaptation (e.g., for loops), figure cleaning (e.g., axis ordering), and general error debugging. Figure panel construction and minor adjustments to figures (e.g., text size) were performed using Adobe Illustrator.

## Data Availability

Proteomic data, raw mass spectrometry files, and databases used are deposited in the ProteomeXchange Consortium via the PRIDE partner repository (Deutsch et al., 2023; Perez-Riverol et al., 2022) and will be publicly available prior to publication and access can be provided upon request. See **Supplementary Data 12** for normalized data used in analyses.

## Supplementary Information

**Supplementary Data 1.** Summary statistics of LC-MS/MS run and database search across study.

**Supplementary Data 2.** Results of compositional statistics including permutational ANOVA via adonis2() and permutational multivariate dispersion analysis via betadisper() and permutest() in vegan (2.6-4) on square root transformed TSS AUC data.

**Supplementary Data 3.** Differential expression analysis via Welch T-test with Benjamini-Hochberg FDR correction showing enrichment of proteins in each proteomic fraction compared the remaining two fractions.

**Supplementary Data 4.** GO Enrichment of enriched proteins in each fraction compared to the bulk proteome by over representation analysis using ClusterProfiler.

**Supplementary Data 5.** Differential expression analysis at the protein level of symbiont carrying versus symbiont free wasps using MaAsLin2.

**Supplementary Data 6.** Differential expression analysis at the orthogroup level of symbiont carrying versus symbiont free wasps using MaAsLin2.

**Supplementary Data 7.** Key term search approach used for identifying immune and odorant binding proteins from annotation data.

**Supplementary Data 8.** Curated immune system annotation based on key term search results.

**Supplementary Data 9.** *Encarsia* proteins that respond to symbiont infection that may be involved in cellular signaling, cell cycle regulation, and/or cell division.

**Supplementary Data10.** Curated odorant binding protein annotation based on key term search results.

**Supplementary Data 11.** Protein annotation data for the *Encarsia suzannae* proteome.

**Supplementary Data 12.** Total sum scaled (TSS) area under the curve (AUC) quantified *Encarsia suzannae* proteome used for analyses in this study.

## Author Contributions

Conceptualization, O.L.M, M.S.H., and M.K.; Methodology, O.L.M., S.E.K., M.R.D., D.L.S., S.S., M.S.H., and M.K.; investigation, O.L.M., S.E.K.; formal analysis, O.L.M.; writing - original draft, O.L.M; writing - review & editing, O.L.M, S.E.K., M.R.D., D.L.S., S.S., M.S.H., and M.K.; visualization - O.L.M.; supervision, M.R.D., M.S.D., S.S., and M.K.; funding acquisition, O.L.M., M.S.H., S.S., and M.K.

## Acknowledgments

We would like to thank Meagan Ash for microscopy and image processing training and discussions, as well as Aram Mikaelyan and Ayesha Awan for thoughtful discussions and feedback on figures. This work was funded by the National Science Foundation awards: #2426305 and #2003107 to M. Kleiner, #2002934 and #2426306 to M.S. Hunter, and #2002987, #2426304, and #2545003 to S. Schmitz-Esser. This work was also supported by The Global One Health Academy at North Carolina State University through a graduate travel award to O.L. Mathieson. We made all LC-MS/MS measurements in the Molecular Education, Technology, and Research Innovation Center (METRIC) at North Carolina State University, which is supported by the State of North Carolina, USA.

