## Supporting Information for "Parasitoid wasp response to infection with a reproductive manipulator symbiont reveals subtle changes in host proteome in life-long symbiosis"

### Supplementary Figures

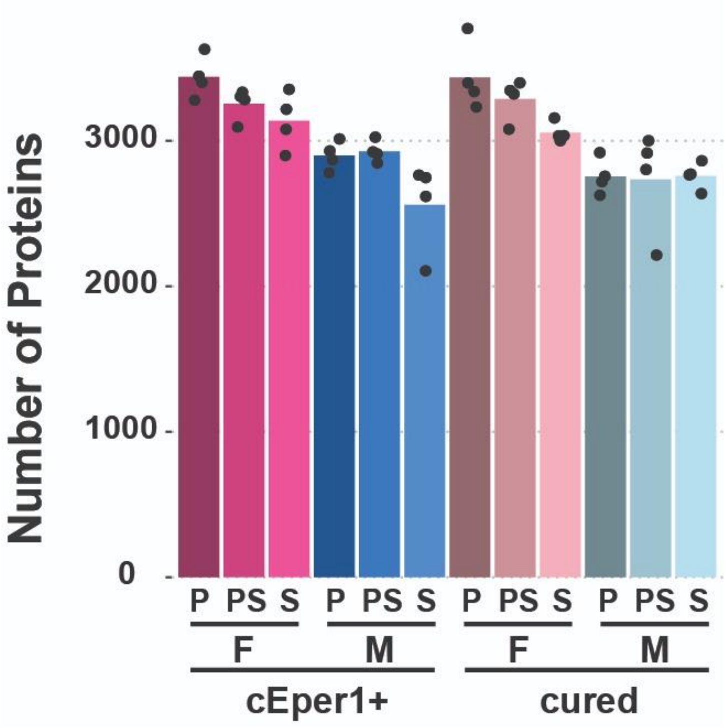


**Figure S1. AUC-quantifiable proteins detected across fraction, sex, and infection status**. Bars indicate mean number of proteins and points indicate the number of proteins in individual samples.

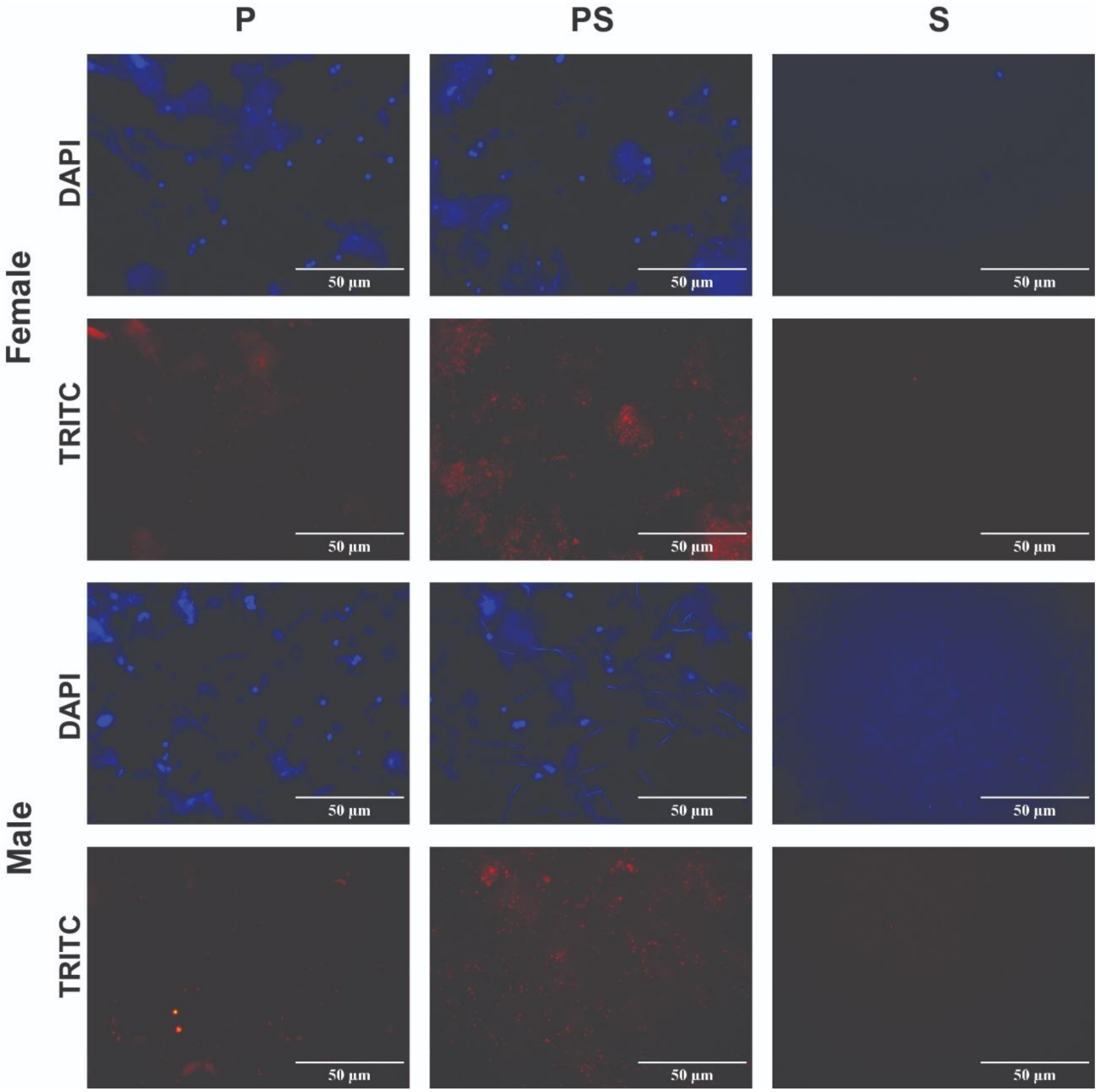


**Figure S2. Fluorescence microscopy of fractionated homogenates.** Blue fluorescence is DAPI DNA stain and red fluorescence is symbiont cy3 probes.

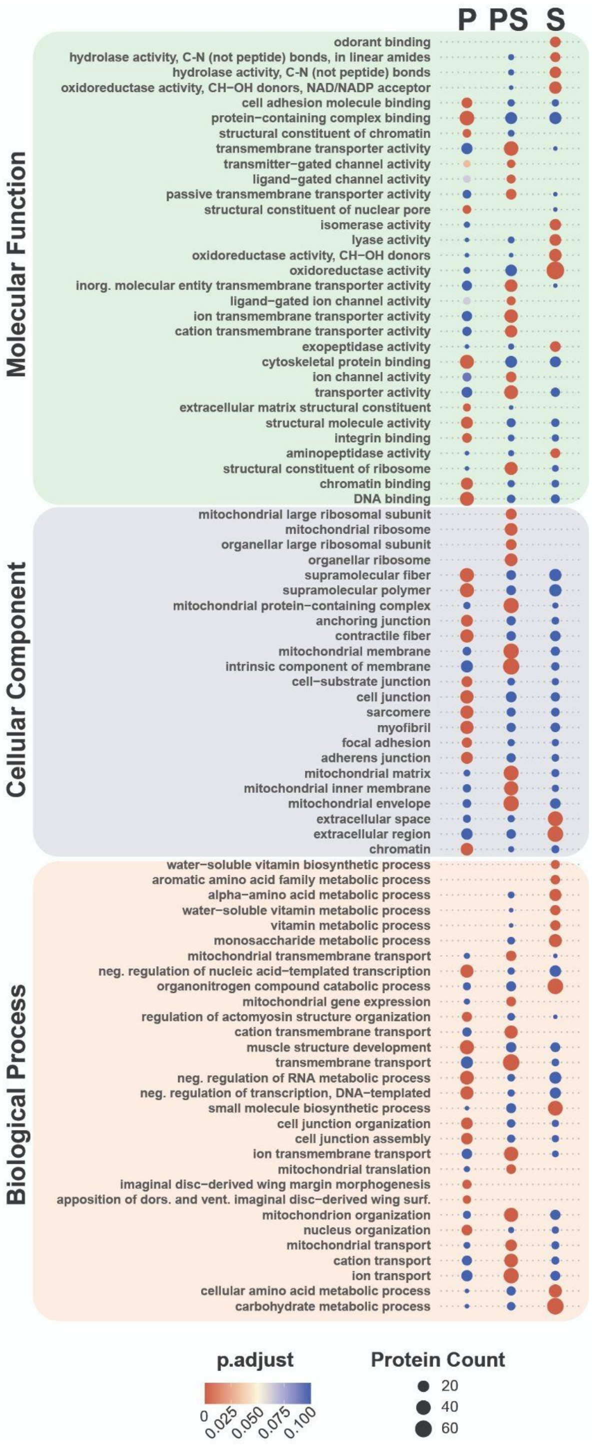


**Figure S3. Enriched GO terms by fraction.** Top significantly enriched GO terms of enriched proteins from each fraction by over representation analysis (enricher function; clusterProfiler R package) compared to the bulk proteome. Significantly enriched proteins were determined by Welch's T-test (see methods). Missing point indicates that there were no proteins significantly enriched in the fraction proteome for a particular GO category, not that the GO category was absent from the fraction proteome.

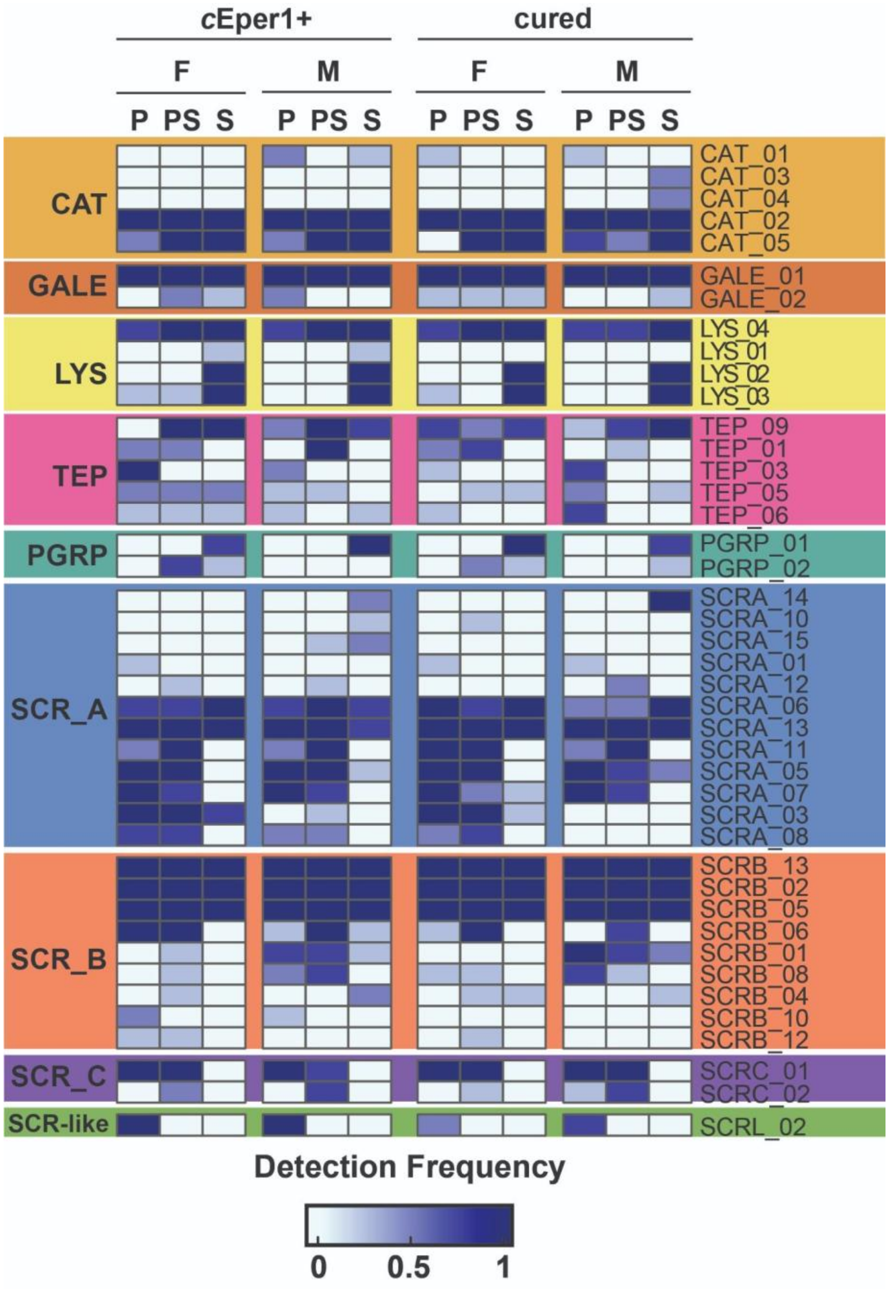


**Supplementary Figure 4. Frequency of detection of immune proteins by category across fraction, sex, and infection status.** Abbreviations: CAT = catalases, GALE = galectins, LYS = lysozymes, PGRP = peptidoglycan recognition proteins, SCR-A (B, C, like) = scavenger receptors class A (B,C, like), TEP = thio-ester containing proteins.

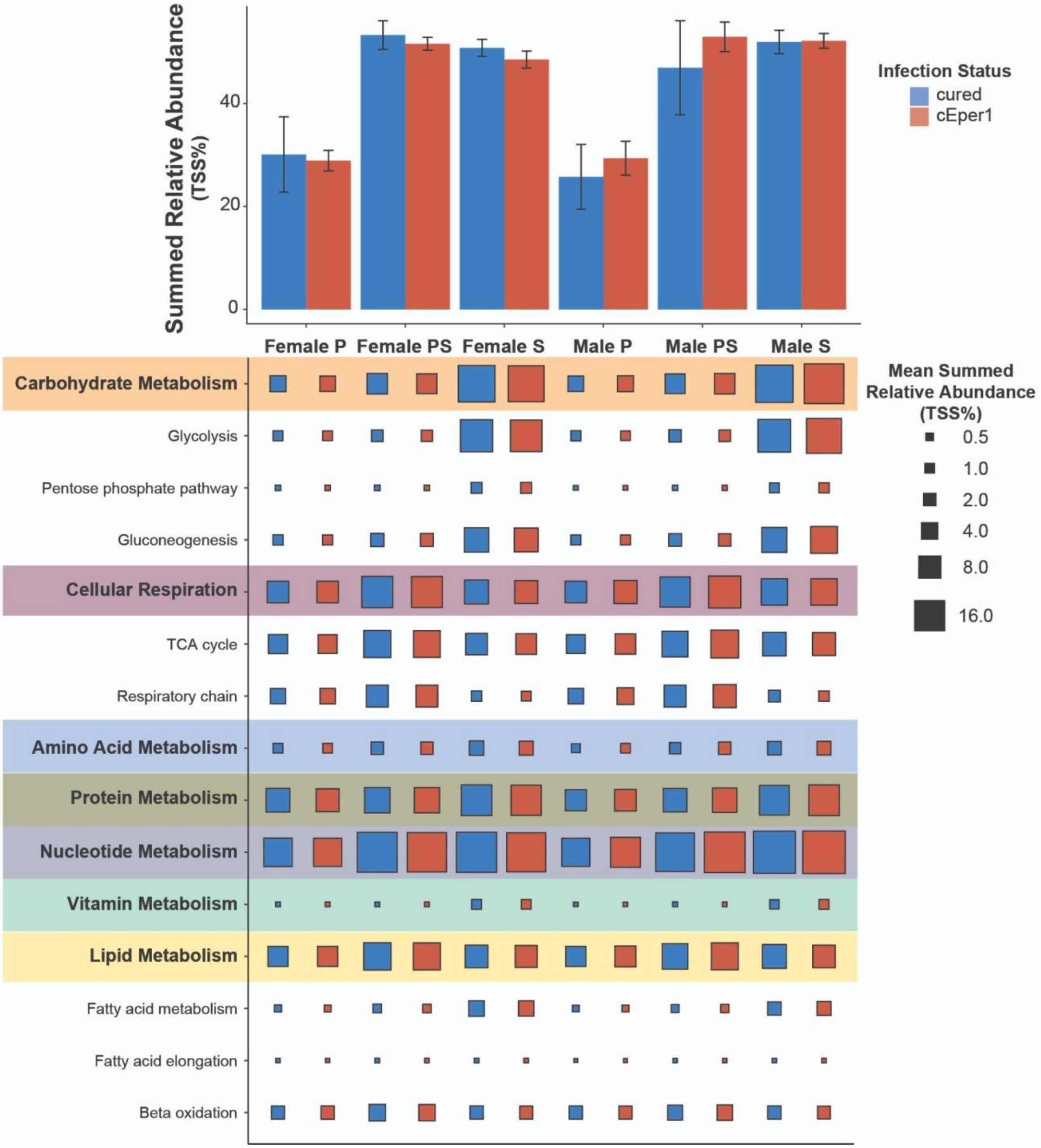


**Supplementary Figure 5. Impact of symbiont infection on metabolism.** Bars indicate average summed abundance of all proteins assigned GO terms descended from "metabolic processes" and error bars indicate 95% confidence intervals. Bubble plot shows average summed abundance of specific subsets of metabolism.

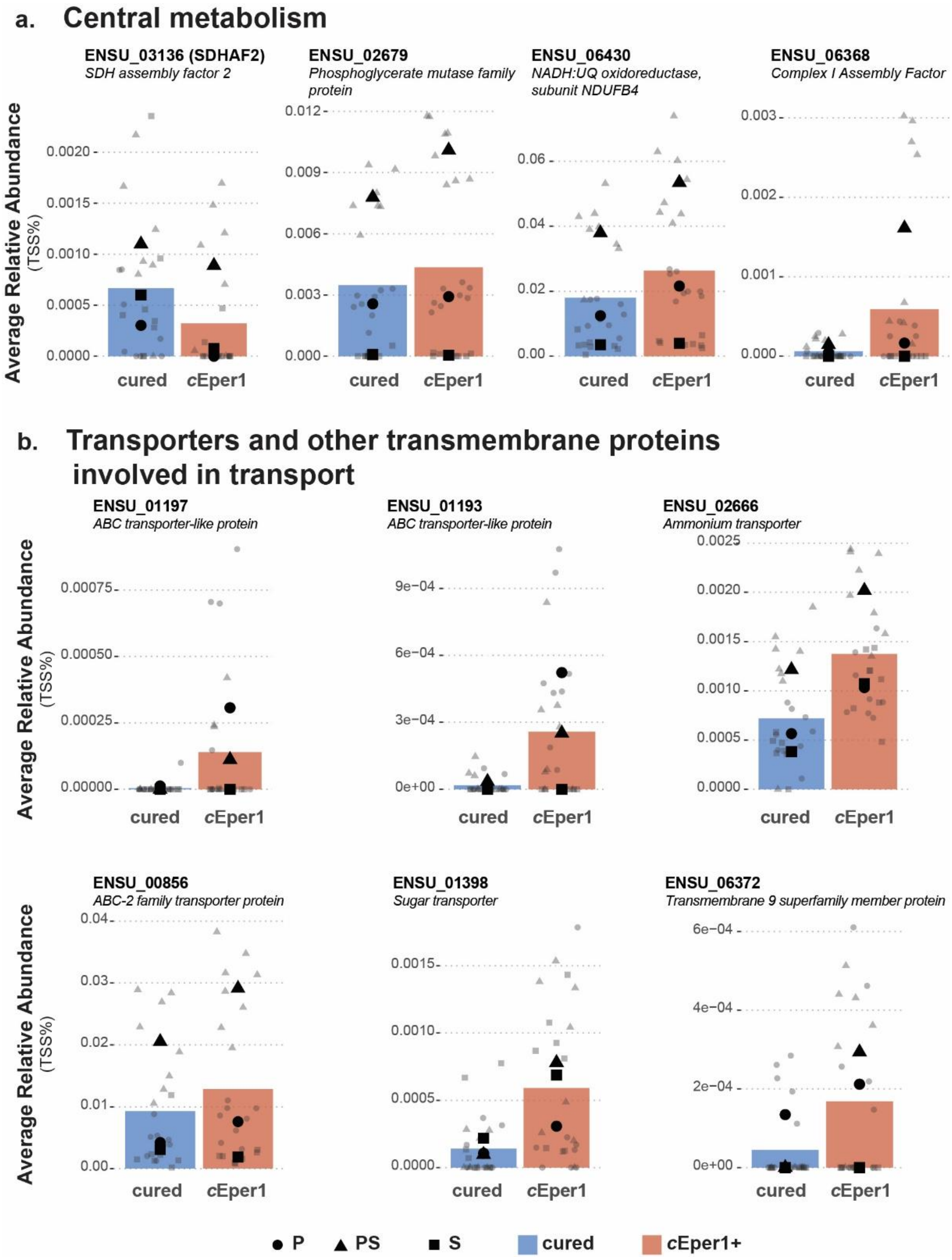


**Supplementary Figure 6. Detection of significantly differentially expressed proteins likely involved in central metabolism and substrate transport.** Bars represent average relative abundance (AUC, TSS%) for a given protein by infection status and individual points represent relative abundance per replicate. The shape indicates proteome fraction, and larger black shapes indicate the average of the fraction (circle = P fraction, triangle = PS fraction, square = S fraction). Colors: blue = cured, orange = *c*Eper1+.

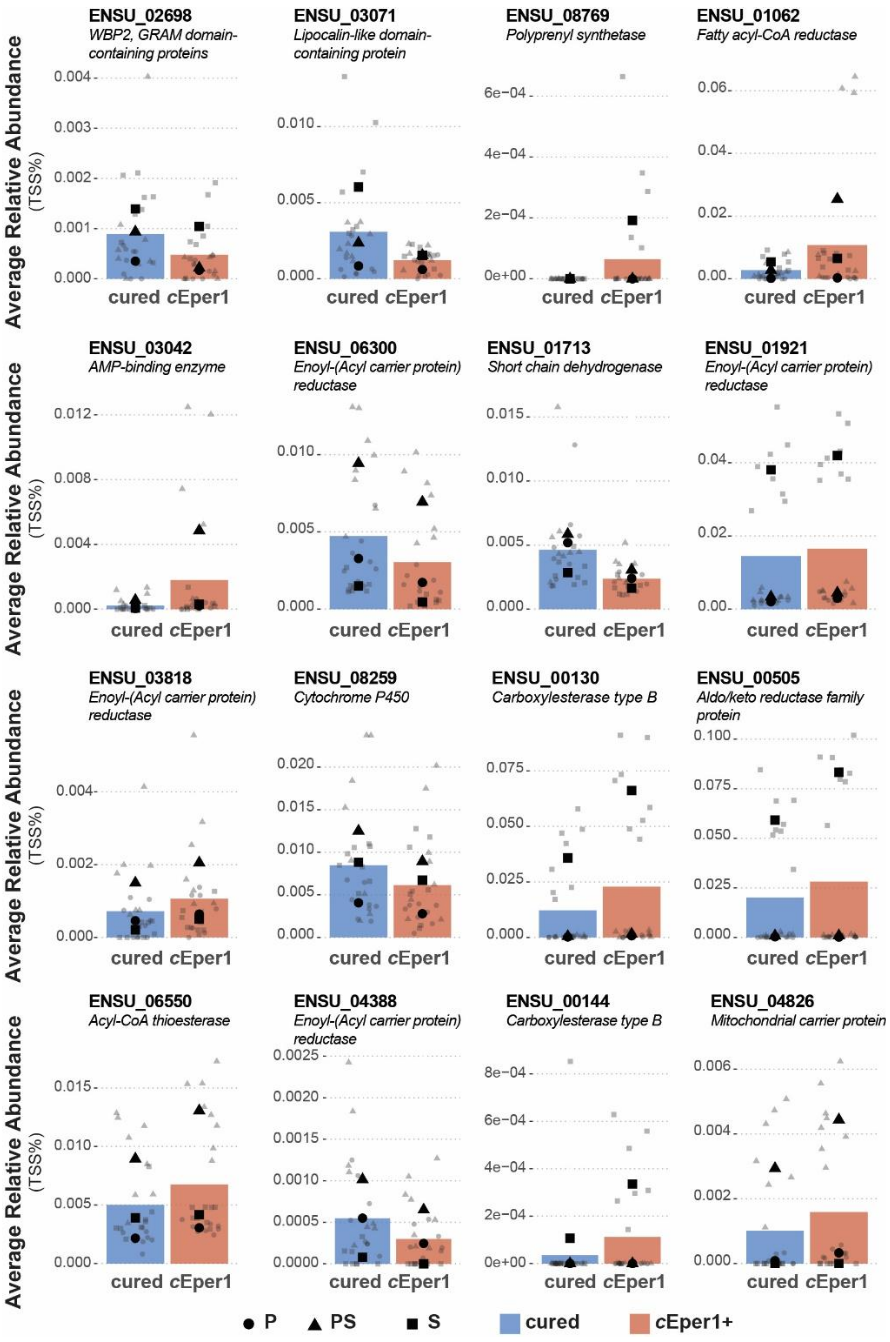


**Supplementary Figure 7**. **Detection of significantly differentially expressed proteins likely involved in fatty acid and lipid metabolism.** Bars represent average relative abundance (AUC, TSS%) for a given protein by infection status and individual points represent relative abundance per replicate. The shape indicates proteome fraction, and larger black shapes indicate the average of the fraction (circle = P fraction, triangle = PS fraction, square = S fraction). Colors: blue = cured, orange = *c*Eper1+.

### Supplementary Table

**Supplementary Table 1. Results of compositional statistics.** Tests were performed using vegan (v2.6-4) adonis2() (PERMANOVA) and betadisper + permutest (PERMDISP) on square root transformed TSS AUC data.

**Protein-level Composition**

| | **Weighted** | | **Unweighted** | |
|---|---|---|---|---|
| | **PERMANOVA** | **PERMDISP** | **PERMANOVA** | **PERMDISP** |
| | ***p-value*** | ***p-value*** | ***p-value*** | ***p-value*** |
| **Fraction** | **0.001** | **0.545** | **0.001** | **0.671** |
| **Sex** | **0.005** | **0.701** | **0.004** | **0.103** |
| **Infection** | **0.957** | **0.933** | **0.982** | **0.808** |
| **Type** | **0.001** | **0.077** | **0.001** | **0.035** |

**Homolog-level Composition**

| | **Weighted** | | **Unweighted** | |
|---|---|---|---|---|
| | **PERMANOVA** | **PERMDISP** | **PERMANOVA** | **PERMDISP** |
| | ***p-value*** | ***p-value*** | ***p-value*** | ***p-value*** |
| **Fraction** | **0.665** | **0.202** | **0.683** | **0.261** |
| **Sex** | **0.644** | **0.332** | **0.709** | **0.911** |
| **Infection** | **0.057** | **0.653** | **0.057** | **0.276** |
| **Type** | **0.88** | **0.932** | **0.883** | **0.860** |
